# Newly synthesized mRNA escapes translational repression during the acute phase of the mammalian unfolded protein response

**DOI:** 10.1101/2022.03.06.483204

**Authors:** Mohammed R. Alzahrani, Bo-Jhih Guan, Leah L. Zagore, Jing Wu, Donny D. Licatalosi, Kristian E. Baker, Maria Hatzoglou

**Affiliations:** Department of Biochemistry, Case Western Reserve University, Cleveland, OH 44106, USA; Department of Genetics and Genome Sciences, Case Western Reserve University, Cleveland, OH, 44106, USA; Center for RNA Science and Therapeutics, Case Western Reserve University, Cleveland, OH 44106, USA

**Keywords:** Poly(A) Tail Length, Translation Inhibition, mRNA Stability, ER Stress, UPR, XBP1

## Abstract

Endoplasmic Reticulum (ER) stress, caused by the accumulation of misfolded proteins in the ER, elicits a homeostatic mechanism known as the Unfolded Protein Response (UPR). The UPR reprograms gene expression to promote adaptation to chronic ER stress. The UPR comprises an acute phase involving inhibition of bulk protein synthesis and a chronic phase of transcriptional induction coupled with the partial recovery of protein synthesis. However, the role of transcriptional regulation in the acute phase of the UPR is not well understood. Here we analyzed the fate of newly synthesized mRNA encoding the protective and homeostatic transcription factor X-box binding protein 1 (XBP1) during this acute phase. We have previously shown that global translational repression induced by the acute UPR was characterized by decreased translation and increased stability of *XBP1* mRNA. We demonstrate here that this stabilization is independent of new transcription. In contrast, we show *XBP1* mRNA newly synthesized during the acute phase accumulates with long poly(A) tails and escapes translational repression. Inhibition of nascent RNA polyadenylation during the acute phase decreased cell survival with no effect in unstressed cells. Furthermore, during the chronic phase of the UPR, levels of *XBP1* mRNA with long poly(A) tails decreased in a manner consistent with co-translational deadenylation. Finally, additional pro-survival, transcriptionally-induced mRNAs show similar regulation, supporting the broad significance of the pre-steady state UPR in translational control during ER stress. We conclude that the biphasic regulation of poly(A) tail length during the UPR represents a previously unrecognized pro-survival mechanism of mammalian gene regulation.

## Introduction

Folding and processing of native polypeptides, the primary function of the Endoplasmic Reticulum (ER), are mediated by a cadre of ER-resident protein chaperones. ER dysfunction leads to an increase in misfolded proteins in the ER lumen, a condition known as ER stress (Adams et al. 2020). ER stress activates ER-transmembrane proteins, including Protein RNA-like Endoplasmic Reticulum Kinase (PERK), Inositol-Regulated Enzyme 1-alpha (IRE1α), and Activating-Transcription Factor 6-α (ATF6α). Together, these proteins sense the misfolded proteins in the ER and activate a protein quality control pathway known as the Unfolded Protein Response (UPR). The UPR, in turn, reprograms gene expression at both transcriptional and translational levels to alleviate cell damage (Guan et al. 2017). The UPR is initiated with an acute phase involving reprogramming of translation, which severely attenuates global protein synthesis while promoting the translation of a select subset of pro-survival mRNAs. Upon sustained stress, both transcriptional and translational reprogramming occurs to coordinate the adaptation to chronic ER stress conditions (Hetz and Papa 2018). Although global translational inhibition during the acute phase of the UPR is partially restored in the late response, translational regulation remains a significant component of adaptation to chronic ER stress (Guan et al. 2017).

In mammalian cells, activation of PERK, ATF6α, and IRE1α upon accumulation of unfolded proteins is achieved through their dissociation from the ER-resident chaperone, BiP (Binding-Immunoglobulin Protein; HSPA5) (Adams et al. 2019). Importantly, PERK encodes a cytoplasmic kinase that phosphorylates the α subunit of the eukaryotic translation initiation factor 2 (eIF2α) which, in turn, binds eukaryotic initiation factor 2B (eIF2B) and prevents recycling of the active eIF2 complex required for translation initiation (Costa-Mattioli and Walter 2020; Baird and Wek 2012; Wek 2018; Gordiyenko et al. 2019). While the decreased availability of eIF2-containing active ternary complexes results in inhibition of bulk protein synthesis, select mRNAs escape this repression and are translationally upregulated via mechanisms involving upstream open reading frames (uORFs) in their 5’-untranslated regions (UTRs) (Pavitt and Ron 2012; Oyadomari and Mori 2004). Among the transcripts translated during the acute UPR is the *ATF4* mRNA, which encodes a transcription factor considered to be the master regulator of the transcriptional reprogramming that occurs during the UPR (Vattem and Wek 2004; Wortel et al. 2017). ATF6α, a second ER transmembrane protein, is activated in response to ER stress through translocation to the Golgi apparatus and cleavage of its N-terminus by Golgi-resident proteases (Correll et al. 2019). Processed ATF6α has been shown to be a potent transcription factor and is also critical for the UPR (Walter et al. 2018). Finally, IRE1α, a dual-activity protein harboring both cytoplasmic kinase and endoribonuclease (RNase) domains, is activated upon initiation of the UPR through oligomerization (Bashir et al. 2021; Ferri et al. 2020) and conformational changes in the RNase domain that catalyze the removal of a 26-nucleotide intron in the coding sequence of X-box binding protein 1 (*XBP1u*) mRNA (Hetz et al. 2015). This process of splicing is unconventional since it occurs in the cytoplasm in proximity to the ER membrane (Fig. 1A) (Yanagitani et al. 2009).

**Figure 1.**
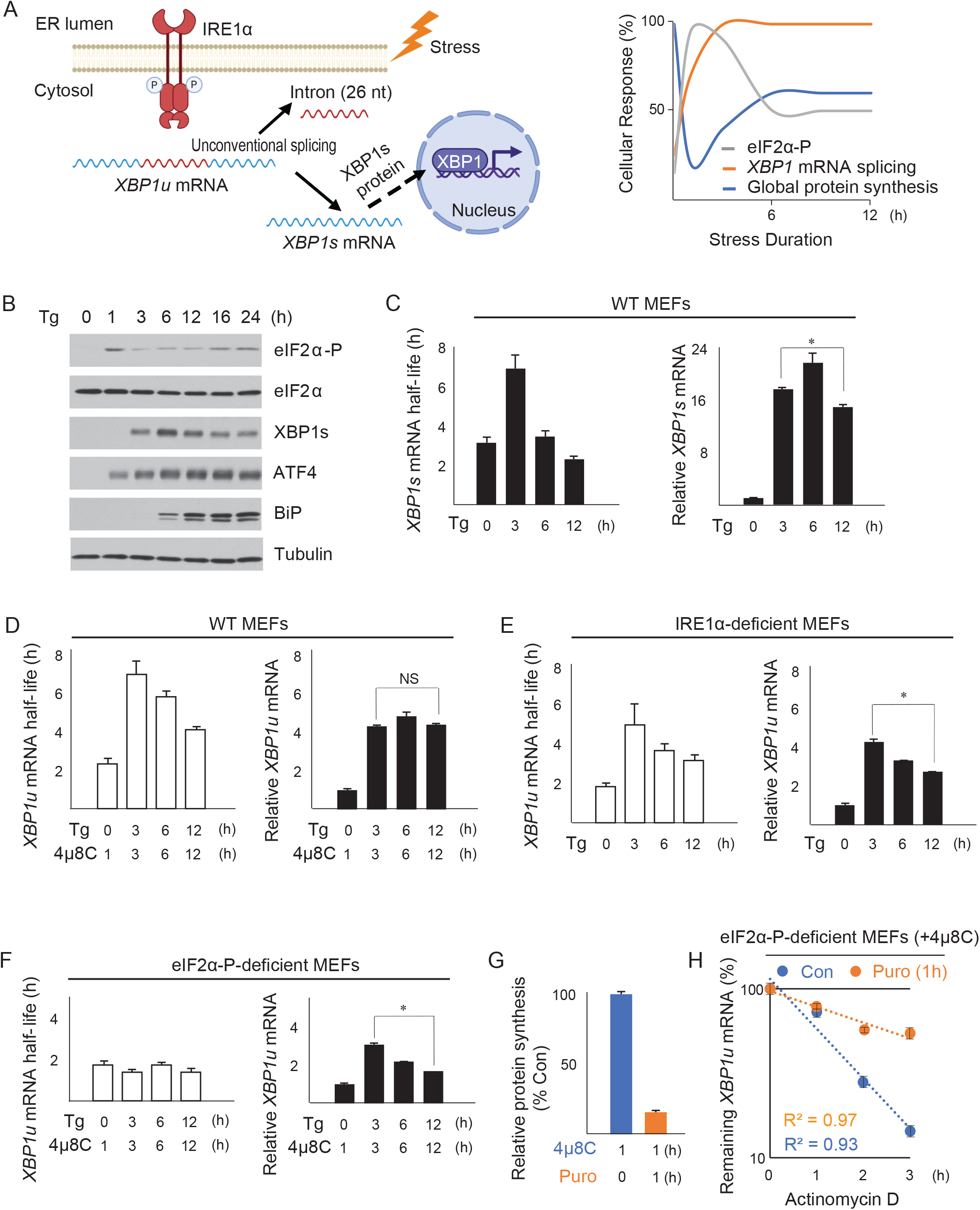
ER stress-induced translation inhibition stabilizes *XBP1u* mRNA during the acute UPR. (A) (Left) Unconventional splicing of *XBP1* mRNA in the cytoplasm. Upon accumulation of misfolded proteins, the cytoplasmic endoribonuclease IRE1α is activated and cleaves a 26-nucleotide (nt) intron from the coding region of the *XBP1u* mRNA. This process occurs in close proximity to the ER membrane. The resulting *XBP1s* mRNA is translated in the cytoplasm, and the XBP1s protein migrates to the nucleus to induce transcription of genes that protect cells from ER stress. The figure was created by Biorender.com (Right) Schematic of temporal responses to ER stress, including eIF2α phosphorylation (eIF2α-P), *XBP1* mRNA splicing, and global protein synthesis rates. (B) Western blot analysis of the indicated proteins in MEFs treated with Tg (400 nM) for the indicated durations. (C) (Left) The half-life of the spliced *XBP1* mRNA (*XBP1s*) was determined by treating MEFs with Tg for the indicated durations. Actinomycin D (10 µg/ml) was added for 0, 0.5, 1, 2, 3, and 4 h at each indicated time of Tg-treatment to inhibit transcription and measure the mRNA decay rate at different time points of ER stress. (Right) The *XBP1s* mRNA levels were quantified by RT-qPCR. (D) The half-life and levels of *XBP1u* mRNA were assessed as in C, except MEF cells were treated with Tg in the presence of a selective inhibitor of the IRE1α endoribonuclease, 4µ8C (50 µM), for the indicated durations. In the absence of Tg treatment in the control condition, MEFs were treated with only 4µ8C for 1 h. (E) The half-life and levels of *XBP1u* mRNA were assessed in MEFs deficient in IRE1α protein (IRE1α-deficient MEFs). The evaluation of half-life and levels of *XBP1u* mRNA was performed as in C. (F) The half-life and levels of the *XBP1u* mRNA were determined in eIF2α-P-deficient MEFs in the presence of 4µ8C as described in C. (G) Protein synthesis was measured by [^35^S]-Met/Cys metabolic labeling in eIF2α-P-deficient MEFs in the presence of 4µ8C alone (Con) or together with Puromycin (Puro, 5 µg/ml) for 1 h to inhibit translation elongation. (H) The half-life of the *XBP1u* mRNA was measured in eIF2α-P-deficient MEFs treated with either 4µ8C for 1 h or Puro and 4µ8C for 1 h. Actinomycin D was added at the indicated times in the presence of 4µ8C or Puro and 4µ8C (t1/2 = Con: 1.2 +/- 0.2 h, Puro: 3.5 +/-0.5 h). Bars represent the mean of 3 independent determinations +/- SEM.

Transcriptional activation upon initiation of the UPR occurs when ATF4, XBP1 (expressed from spliced *XBP1s* mRNA), and ATF6α translocate into the nucleus and induce expression of target genes that establish the adaptive phase of UPR that restores cellular proteostasis during chronic ER stress (Hetz 2012). Notably, transcription of *XBP1* is regulated by both XBP1 and ATF6, suggesting intersection of the two separate pathways of the UPR and highlighting the importance of the expression of XBP1 during the UPR (Park et al. 2021; Tsuru et al. 2016; Sharma et al. 2020). The target genes of the transcription program initiated by these three UPR effectors include additional transcription factors, which may control the threshold of transcriptional activation of target genes; this is important given that excessive induction of UPR target genes can be harmful during chronic ER stress, especially when global protein synthesis rates are not well balanced with the capacity of the ER (Oslowski and Urano 2011; Spaan et al. 2019; Bartoszewska and Collawn 2020; Han et al. 2013; Wek 2018). Indeed, negative feedback mechanisms limiting transcriptional reprogramming during UPR, have been described (Li et al. 2020; Amin-Wetzel et al. 2017; Oslowski and Urano 2011; Bartoszewski et al. 2008; Gonen et al. 2019), including induction of transcriptional repressors (Ku and Cheng 2020) and microRNA-mediated regulation of specific mRNA targets (Zhao et al. 2020; Maurel and Chevet 2013).

Transition in the cell from the acute to chronic UPR is mediated by a network of integrated signaling pathways (Adams et al. 2019; You et al. 2021; Guan et al. 2017; Hetz et al. 2015). The mechanisms of translational control during the progression to adaptation involve signaling downstream of activation of the PERK kinase (Harding et al. 2000). ER stress-activated PERK attenuates eIF4E-mediated mRNA translation, thus limiting translation of most mRNAs in the chronic phase. At the same time, the decreased eIF2B activity downstream of eIF2 phosphorylation during both acute and chronic ER stress sustains the selective translation of mRNAs with specific 5’-UTR features (Guan et al. 2017; Wek 2018). Among the post-transcriptional mechanisms required for transitioning to chronic UPR is the regulation of mRNA degradation. Genome-wide studies in HeLa cells exposed to chemical-induced ER stress revealed mRNAs with both increased and decreased turnover rates for translationally-induced and translationally-repressed transcripts, suggesting that a subset of mRNAs are stabilized despite being translationally repressed (Kawai et al. 2004). Indeed, we have previously shown that stabilization of *XBP1s* mRNA is coupled to its translational repression during the acute UPR, and during chronic UPR, *XBP1* transcripts become destabilized (Majumder et al. 2012). In contrast, poly(A) tail-end sequencing (TED-seq) during ER stress of HEK293 cells exposed to a chemical stressor suggested a correlation between long poly(A) tail mRNA and increased stability and translation for a number of stress-induced mRNAs (Woo et al. 2018). Among these mRNAs was the *XBP1* mRNA that was found to be less translationally repressed compared to other mRNAs. The discrepancy in the findings between these two studies could be attributed to differences in regulation between *XBP1s* and *XBP1u* mRNAs and the interplay between transcriptional activation and global translational repression during the UPR or may reflect the timing of ER stress (acute or chronic phase of the UPR) investigated in these two studies. To address this unresolved question, we dissected the interplay between transcriptional control and both translational efficiency and mRNA stability in a temporal manner during the UPR, for both *XBP1s* and *XBP1u* mRNAs.

While transcriptional upregulation has been studied under a number of stress conditions (Guan et al. 2017; Gonen et al. 2019; Mahat et al. 2016; Shih et al. 2021), the interplay between transcriptional activation, translation status, and turnover of the resulting transcripts during the acute and chronic phases of the mammalian UPR is less understood. Recent advances in nascent RNA metabolic labeling methods in studies of mRNA synthesis and degradation dynamics (Rabani et al. 2011; Battich et al. 2020; Eisen et al. 2020) now allow for the analysis of the translation and turnover of newly synthesized, stress-induced mRNAs during the progression from the acute to chronic UPR, which can be compared to global translation and mRNA turnover to reveal differences in the regulatory programs of those pools of mRNAs. Such studies can reveal the importance of early transcriptional activation in a cell subjected to global translational repression.

In this study, we hypothesized that the acute UPR phase might include mechanisms to protect stress-induced mRNAs from translational repression. Such cellular mechanisms would be valuable for limiting commitment to the sophisticated and energy-demanding transcriptional and translational reprogramming of the chronic UPR in anticipation of recovery from stress following the acute phase (Walter et al. 2015; Walter and Ron 2011; Corazzari et al. 2017 Krokowski et al. 2013; Han et al. 2013; Wang and Kaufman 2016). Herein we investigated *XBP1* mRNA as a model to understand temporal gene regulation during the acute and chronic UPR phases in mammalian cells. We found that the transcriptional induction of the *XBP1* gene during acute stress and the short half-life of the *XBP1* mRNA results in the rapid substitution of pre-stress *XBP1* mRNAs for newly synthesized mRNAs in cells transitioning from the acute to chronic UPR. We also show that newly synthesized *XBP1* mRNA escapes translational repression during the acute UPR but is rapidly turned over during the chronic UPR. In addition, we show *XBP1* mRNA poly(A) tail length is dynamic, with transcripts harboring long poly(A) tails during the acute UPR rapidly transitioning to a short poly(A) tail during the chronic UPR. In contrast to the regulation of the newly synthesized *XBP1* mRNA, we previously showed and confirmed here that *XBP1s* mRNA is translationally repressed and stabilized during the acute UPR (Majumder et al. 2012). This mechanism likely involves coupled translational repression and decreased turnover of *XBP1* mRNA and is independent of new transcription during the acute UPR. On the other hand, increased mRNA stability of the translationally repressed mRNA during the acute UPR, allows rapid translation and *XBP1s* protein accumulation during recovery from acute ER stress. We conclude that long poly(A) tails in newly synthesized mRNA during the pre-steady state UPR facilitate escape from translational repression, whereas co-translational shortening of the poly(A) tails during chronic ER stress limits translation and stability of the mRNAs, leading to a decreased threshold of the UPR to homeostatic levels. Our analysis of *XBP1* mRNA reveals a previously unrecognized mechanism of gene regulation dependent upon mRNA poly(A) tail length and illuminates how transcriptional and post-transcriptional programs are coupled to mediate the cellular response to stress.

## Results

### Acute UPR-induced translation inhibition stabilizes *XBP1u* mRNA

To study the regulation of the *XBP1* gene during the UPR, we treated mouse embryonic fibroblasts (MEFs) with Thapsigargin (Tg), an irreversible inhibitor of the Sarcoplasmic/Endoplasmic Reticulum Calcium ATPase (SERCA) pump that leads to accumulation of unfolded proteins in the ER (Sehgal et al. 2017). We first determined that this experimental system elicits the expected temporal regulation of *XBP1* gene expression as cells transition from acute to chronic UPR. Temporal changes in protein synthesis upon Tg-treatment are caused by the transient increase in phosphorylation of eukaryotic translation initiation factor eIF2α (Fig. 1A and 1B) (Dorrbaum et al. 2020; Rothenberg et al. 2018). Although splicing of the *XBP1u* mRNA can occur rapidly in response to ER stress (Supplemental Fig. S1A) (Tam et al. 2014), accumulation of XBP1s protein is delayed (Fig. 1B). In contrast, ATF4 is rapidly induced during the acute UPR, and BiP during the transitioning to the chronic UPR (Fig. 1B) (Gonen et al. 2019; Guan et al. 2017). The delayed accumulation of XBP1s protein is in agreement with our earlier findings that *XBP1s* mRNA is translationally repressed during the acute UPR and translationally de-repressed during the chronic UPR (Majumder et al. 2012). Furthermore, we confirmed our earlier report that *XBP1s* mRNA is stabilized during the acute UPR and destabilized during the chronic UPR in MEFs (Fig. 1C). Notably, the UPR-induced transcription of the *XBP1* gene contributed to a several-fold increase in *XBP1s* mRNA in the acute UPR phase (Fig. 1C) (He et al. 2010).

To better understand the role of transcriptional and post-transcriptional regulation of the *XBP1* gene during the UPR, we investigated if regulation of *XBP1u* mRNA is similar to *XBP1s* mRNA. We measured the half-life and abundance of *XBP1u* mRNA in MEFs during the UPR using 4μ8C, a pharmacological inhibitor of IRE1α (Supplemental Fig. S1A) (Cross et al. 2012). Similar to *XBP1s* mRNA, *XBP1u* mRNA was found to be translationally repressed (M Alzahrani, unpublished data) and transiently stabilized during the acute UPR (Fig. 1D). Notably, levels of *XBP1u* mRNA also showed an increase during the acute phase of UPR (Fig. 1D). We next used IRE1α-deficient MEFs lacking IRE1α splicing activity (Lee et al. 2002) to monitor *XBP1u* mRNA levels during UPR (Supplemental Fig. S1A). We found that regulation of *XBP1u* mRNA was indistinguishable from 4µ8C-treated MEFs as such that during the acute UPR, *XBP1u* mRNA was stabilized and its level increased, followed by destabilization and a decrease in transcript abundance during the chronic UPR (Fig. 1E). Similar to *XBP1s* mRNA (Majumder et al. 2012), translation of the *XBP1u* mRNA was repressed in the acute UPR and de-repressed in the chronic phase in these cells (Supplemental Fig. S2A). As anticipated, translation of the *ATF4* mRNA increased in the acute phase and remained high in the chronic phase, confirming the experimental system (Fig. 1B and Supplemental Fig. S2B). These data support that regulation of *XBP1u* mRNA stability and translation during the UPR is similar to regulation of the *XBP1s* mRNA.

To address if translational reprogramming mediated by eIF2α phosphorylation during the acute UPR contributes to increased stability of *XBP1u* mRNA, we monitored *XBP1u* mRNA in MEFs that express a mutant eIF2α unable to be phosphorylated at serine 51 (eIF2α-P-deficient MEFs). In these cells, eIF2α is not responsive to the UPR and, consequently, translation initiation is not attenuated (Scheuner et al. 2001). We found that despite increased abundance of the *XBP1u* mRNA, transcripts were not stabilized during the acute UPR in these cells (Fig. 1F). Importantly, translation of *XBP1u* mRNA was not inhibited during the acute UPR phase in eIF2α-P-deficient MEFs, as expected (Supplemental Fig. S2C). Together, these data indicate that *XBP1s* and *XBP1u* mRNAs are subject to similar regulation during the UPR, and that translational repression mediated by eIF2α phosphorylation is critical for the increased stability of the *XBP1* mRNA in the acute phase of the stress response. Because the increased levels of the *XBP1u* mRNA in eIF2α-P-deficient cells was the result of transcriptional activation and not increased mRNA stability, we also conclude that stabilization of the *XBP1* mRNA during the acute UPR phase is not coupled to transcriptional induction of the *XBP1* gene (Fig. 1F), as has been previously suggested (Slobodin et al. 2020).

To better understand the underlying mechanism contributing to the transient stabilization of the *XBP1* mRNA, we inhibited translation in the absence of ER stress. eIF2α-P-deficient MEFs were treated with puromycin to inhibit translation elongation, and protein synthesis was monitored by metabolic labeling (in the presence of 4µ8C). As expected, cells treated with puromycin for one hour showed a dramatic decrease in global protein synthesis (Fig. 1G). Interestingly, *XBP1u* mRNA half-life increased in puromycin-treated MEFs despite the absence of an acute UPR (Fig. 1H), a result that was recapitulated using harringtonine, an inhibitor of translation initiation (Supplemental Fig. S1B). These data strongly suggest that *XBP1* mRNA levels and stability are tightly correlated with the translation status of the cell and that transcriptional induction of *XBP1* transcription alone is insufficient to lead to the observed increase in *XBP1* mRNA abundance during ER stress.

### The acute phase of UPR does not increase bulk mRNA poly(A) tail length

mRNA poly(A) tails have been proposed to regulate post-transcriptional gene expression at the levels of mRNA translation and stability (Krause et al. 2019). A recent genome-wide analysis has demonstrated a positive correlation between the median poly(A) tail length per transcript and its mRNA half-life in fibroblast cells during steady state conditions (Chang et al. 2014). Little is known, however, about how poly(A) tail length is regulated during progression of the UPR from the acute to chronic phase and the role tail length may play in response to stress. Here, we used the transcriptionally induced *XBP1* mRNA to determine whether a correlation exists between the observed changes in mRNA half-life during the UPR and the length of its poly(A) tail. We employed a PCR-based poly(A) tailing assay to estimate the length of *XBP1* mRNA poly(A) tails under UPR conditions (Patil et al. 2014). We found that the poly(A) tail length of *XBP1* mRNA varied dynamically during the UPR, with the poly(A) tail length being short at steady state before stress (at 0 time) and predominantly long during the acute UPR, followed by a decrease in length during the transition to the chronic UPR. During the acute UPR, two populations of *XBP1* mRNA with either long or short poly(A) tails were detected, with the long poly(A) tail population being the predominant species at 3 h of ER stress (Fig. 2A). The poly(A) tails of the *GAPDH* mRNA, which is not induced in abundance during the UPR, did not change between the untreated and the acute UPR, but became shorter during the transition to the chronic UPR (Fig. 2A).

**Figure 2.**
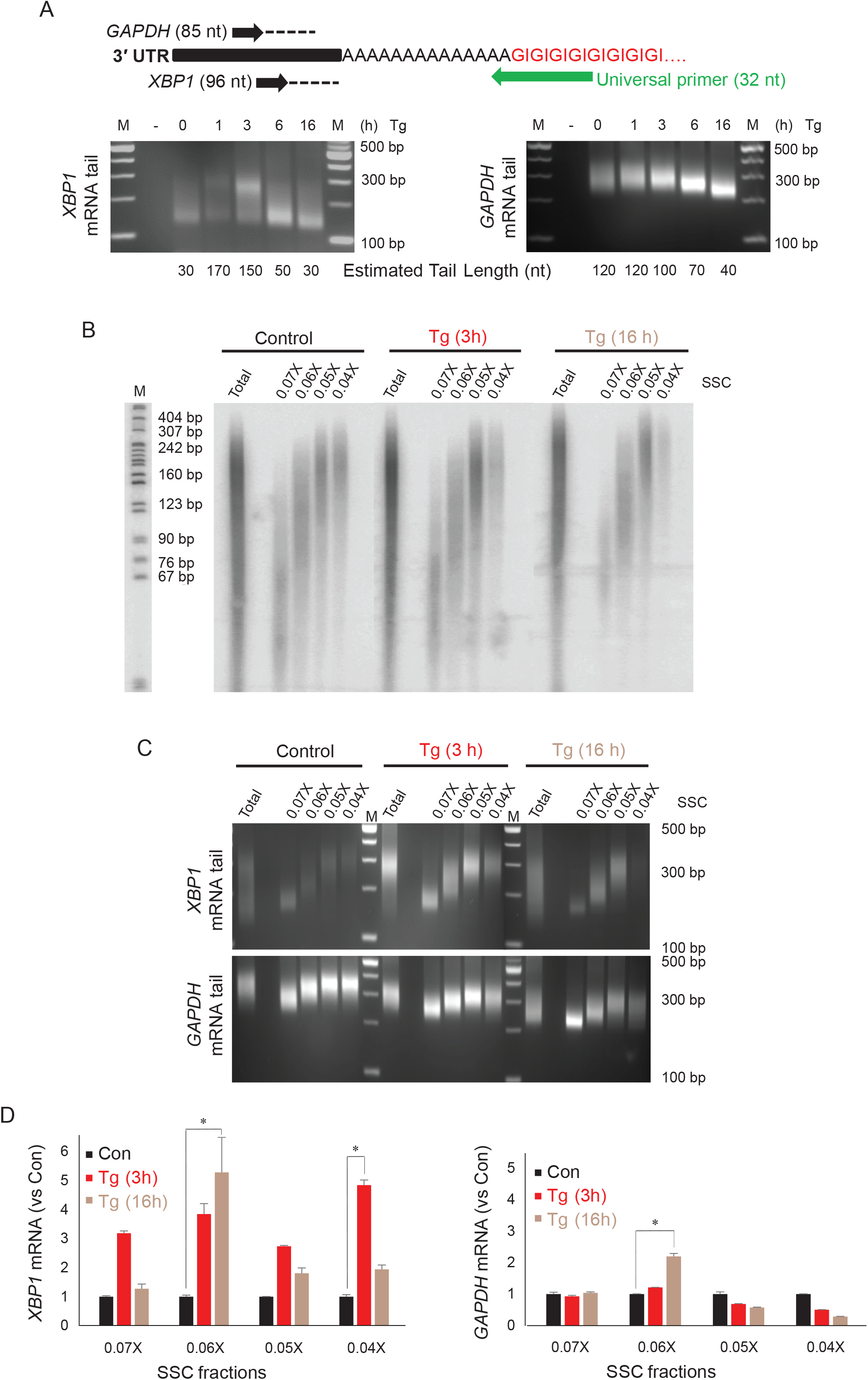
Acute UPR does not involve an increase of global poly(A) tail length of mRNAs. (A) (Top) Experimental diagram of the PCR-based poly(A) tailing assay (Patil et al. 2014) for *XBP1* and *GAPDH* mRNAs. The 3’-end of total RNA was modified with GMP/IMP residues by Poly(A) polymerase to form a GI-oligo tail connected to the poly(A) tail. Following RNA isolation and RT, universal primers were used as a reverse primer for cDNA amplification of targets. (Bottom) PCR-based poly(A) tailing assay in MEFs treated with Tg for the indicated durations. Estimated tail lengths are shown. As a negative control (-), a tail reaction was performed on cDNA derived from RNA not tagged with the GI-oligo tail. (B) mRNA fractionation based on poly(A) tail length in MEFs treated with Tg for the indicated durations. 3 h of Tg treatment represents acute ER stress, and 16 h of Tg treatment represents chronic ER stress. (C) The poly(A) tail length of *XBP1* and *GAPDH* mRNA was estimated using the PCR-based poly(A) tailing assay in the fractionated mRNA pools from B. (D) The level of *XBP1* and *GAPDH* mRNA was determined using RT-qPCR and compared to their own internal controls in all fractionated mRNA pools. Bars represent the mean of 3 independent determinations +/- SEM.

To quantify *XBP1* mRNA harboring different long poly(A) tail lengths in each phase of the UPR, we used two approaches. First, total mRNA from untreated MEFs or cells treated with Tg for 3 h (acute) or 16 h (chronic) was isolated and subjected to fractionation based on the length of mRNA poly(A) tail using decreasing salt concentration from oligo(dT)-bound beads (Meijer and de Moor 2011). Evaluation of the bulk poly(A) tail length distribution in the two phases of the UPR did not show significant changes in mRNA poly(A) tail length in the total mRNA pool as compared to untreated cells (Fig. 2B). In contrast to the bulk poly(A) tail length, there was a sharp increase in mRNAs with long poly(A) tails in cells treated with Tg for 3 h, which was reduced at the 16 h timepoint (Fig. 2A). To identify populations of *XBP1* mRNA harboring different poly(A) tails, we first evaluated poly(A) tail length in eluted fractions using the PCR-based poly(A) tailing assay. We observed increased levels of *XBP1* mRNA with long poly(A) tails in the acute UPR phase compared to the chronic phase of the UPR or in untreated control cells (Fig. 2C). As anticipated, *GAPDH* mRNA with long poly(A) tails was enriched in untreated cells. This enrichment decreased as cells progressed to the acute and chronic UPR phases (Fig. 2C). In a second approach, we performed RT-qPCR analysis for both *XBP1* and *GAPDH* mRNAs in the fractions eluted with different salt concentrations from the oligo(dT) beads, shown in Fig. 2B. We found that during the acute UPR, *XBP1* mRNA with long poly(A) tails showed the highest increase over untreated control (Fig. 2D, 0.04X fraction). In contrast, during the chronic UPR, the highest levels of *XBP1* mRNA was observed to harbor shorter poly(A) tails (Fig. 2D, 0.06X fraction). Notably, both *XBP1* and *GAPDH* mRNA displayed a shortening of their poly(A) tails during the chronic UPR (Fig. 2D, compare *XBP1* Tg (16h) 0.06X fraction to *GAPDH* Tg (16h) 0.06X fraction), suggesting a general mechanism of post-transcriptional regulation as cells adapt to chronic ER stress. These data demonstrate temporal changes in *XBP1* mRNA poly(A) tail length during the UPR and provide a possible mechanism for regulation of *XBP1* gene expression during stress.

### *XBP1* mRNAs with long poly(A) tails escape translational repression during the acute UPR

Global protein synthesis is severely repressed during the acute UPR due to the phosphorylation of the translation initiation factor eIF2α. However, select mRNAs escape translational repression, including the mRNA encoding transcription factor ATF4 (Fig. 1B and Supplemental Fig. S2B) (Vattem and Wek 2004). The predominant mechanism mediating escape from translational repression is through mRNA uORFs in the 5’-UTRs that serve to enhance translation of downstream coding regions when eIF2α is phosphorylated (Wek 2018). However, we and others have found transcripts lacking uORFs associated with ribosomes during the acute UPR (Guan et al. 2017; van den Beucken et al. 2007; Moro et al. 2021; Pavitt and Ron 2012; Jaud et al. 2020). Based on our observation that *XBP1* mRNA is induced and harbors long poly(A) tails, we evaluated whether poly(A) tail length influences the association of the *XBP1* transcript with polyribosomes by polysome profile analysis. We fractionated extracts from MEFs grown under control, acute, and chronic UPR conditions through sucrose gradients (Fig. 3A). As expected, polysome profiles show a dramatic decrease in heavy polysomes and an accumulation of monosomes in Tg-treated cells after 1 h, supporting severe inhibition of translation during the acute UPR. The decreased percentage of monosomes and increased percentage of polyribosomes in Tg-treated cells after 16 h treatment suggests a partial translational recovery during the chronic UPR as previously described (Guan et al. 2017). To evaluate whether poly(A) tail length influences association of the *XBP1* transcripts with ribosomes, we performed the poly(A) tailing PCR-based assay in pooled gradient fractions (Fig. 3A, ribosome-free (F), light polyribosomes (L), and heavy polyribosomes (H)). Strikingly, the majority of long poly(A) tail *XBP1* mRNA was associated with heavy polysomes during the acute UPR, at a time when global translation initiation is compromised (Fig. 3A and 3B). In contrast, we observed a broader range of *XBP1* mRNA poly(A) tail during chronic UPR when inhibition of translation showed a partial recovery (Fig. 3A and 3B). This dynamic change of poly(A) tail length of *XBP1* mRNA on polyribosomes as the cell transitions from the acute to chronic UPR, may reflect co-translational deadenylation during the chronic UPR, as has been previously described in other systems (Duan et al. 2020; Vindry et al. 2012). Taken together, these data suggest that two pools of *XBP1* mRNA exist during the acute UPR: one pool with long poly(A) tails which escapes translational repression and can be found on polyribosomes, and another pool with short poly(A) tails which is translationally attenuated (Fig. 3B). We hypothesize that the long poly(A) tail pool of the *XBP1* mRNA likely represents newly synthesized, stress-induced mRNAs transcribed during the acute UPR. To test this hypothesis, we isolated metabolically labeled, newly synthesized RNA (using 1 h of 5-Ethynyl Uridine (5EU) incorporation) in MEFs grown in control, acute, and chronic UPR conditions, then performed the poly(A) tailing PCR-based assay in the flow-through (old RNA) and eluted (newly synthesized RNA) fractions (Fig. 3C). A homogeneous population of *XBP1* mRNA with long poly(A) tails was enriched in the eluted pool, whereas the flow-through contained *XBP1* mRNA with shorter and heterogeneous poly(A) tail lengths (Fig. 3C). Strikingly, the data demonstrate the newly synthesized *XBP1* mRNA during the acute UPR predominantly harbor long poly(A) tails, in contrast those in the chronic UPR, which showed varied *XBP1* mRNA poly(A) tail length (compare Fig. 3B and 3C). We conclude that newly synthesized *XBP1* mRNA with long poly(A) tails escapes translational repression during the acute UPR. This finding reveals a potential new mechanism for select mRNA translation during the acute UPR dependent upon mRNA poly(A) tail length.

**Figure 3.**
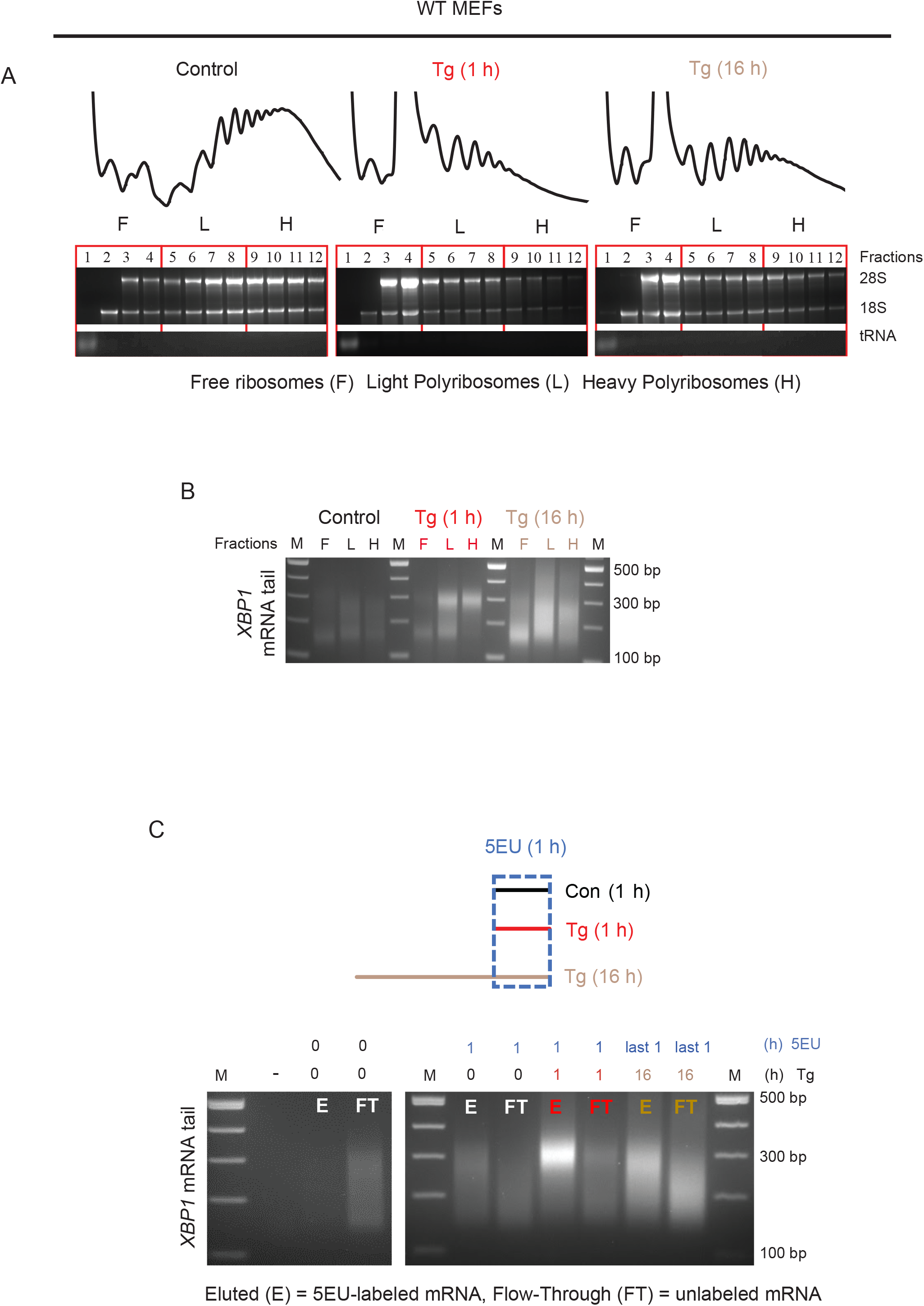
The long tail *XBP1* mRNA escapes translational repression during the acute UPR. (A) Polysome profiles of MEFs treated with Tg for 0, 1 and 16 h. Following RNA isolation and integrity check on agarose gels, RNA was combined into 3 pools as shown; ribosome-free {F}, light polyribosomes {L}, heavy polyribosomes {H}. (B) PCR-based poly(A) tailing assay for *XBP1* mRNAs estimated in the pooled RNA from F, L, or H of Tg-treated MEFs for 0, 1, and 16 h. (C) Experimental schematic and PCR-based poly(A) tailing assay of *XBP1* mRNA was estimated in 5-Ethynyl Uridine (5EU)-labeled RNA isolated from MEFs. Cells were treated with Tg for 1, or 16 h, and 5EU (400 µM) was added for the last 1 h of treatments. Vehicle (DMSO) was added in control cells for 1 h in the presence of 5EU. Following the click chemistry reaction, 5EU-labeled RNA was immobilized by streptavidin beads and was eluted {E}. The flow-through {FT} was also collected as unlabeled RNA. The poly(A) tailing assay was performed on both E and FT pools as previously described and analyzed on denaturing agarose gels. As a negative control, MEFs were not treated with Tg or 5EU. (-) represents a sample of RNA without the GI-tail and serves as a negative control for the data as shown in Figure 2.

### Polyadenylation of nascent transcripts during acute ER stress is critical for cell survival

Although many studies have focused on the importance of transcriptional control in adaptation to chronic ER stress, the importance of acute UPR-induced transcription is less well-studied (Guan et al. 2017; Gonen et al. 2019). To explore the effect of newly synthesized mRNA on cell survival during acute ER stress, we used Cordycepin, an analog of adenosine that blocks the synthesis of RNA with long poly(A) tails on nascent transcripts (Kondrashov et al. 2012), and measured cell viability during the acute UPR. In these experiments, we used Cyclopiazonic Acid (CPA), a reversible inducer of the UPR (Guan et al. 2017), that inhibits the SERCA pump but can be washed out from cells. Similar to Tg treatment, MEFs treated with CPA for 3 h showed a transient accumulation of *XBP1* mRNA with long poly(A) tail during the UPR (compare Figs. 2A and 4A). Importantly, we found no effect of CPA on the viability of MEFs after 3 h (Fig. 4B) and cells after washing out of CPA resumed growth (Fig. 4B). We next inhibited polyadenylation by supplementing the growth media with Cordycepin in the presence or absence of CPA. We found that cell survival was not affected after 3 h in vehicle (DMSO)-, Cordycepin alone-, or CPA-treated MEFs. In contrast, cell viability under the combined treatment of CPA and Cordycepin was significantly compromised (Fig. 4C). Interestingly, Cordycepin treatment decreased *XBP1* mRNA abundance and attenuated the induction of *XBP1* mRNA during the UPR (Fig. 4D). The decreased *XBP1* mRNA abundance was paralleled by decreased accumulation of long poly(A) tail *XBP1* mRNA during the acute UPR (Fig. 4E). Because treatment of MEFs with the transcription inhibitor Actinomycin D during the acute UPR abolished accumulation of *XBP1* mRNA (Fig. 4E), we conclude that Cordycepin predominantly affects polyadenylation of nascent *XBP1* mRNA.

**Figure 4.**
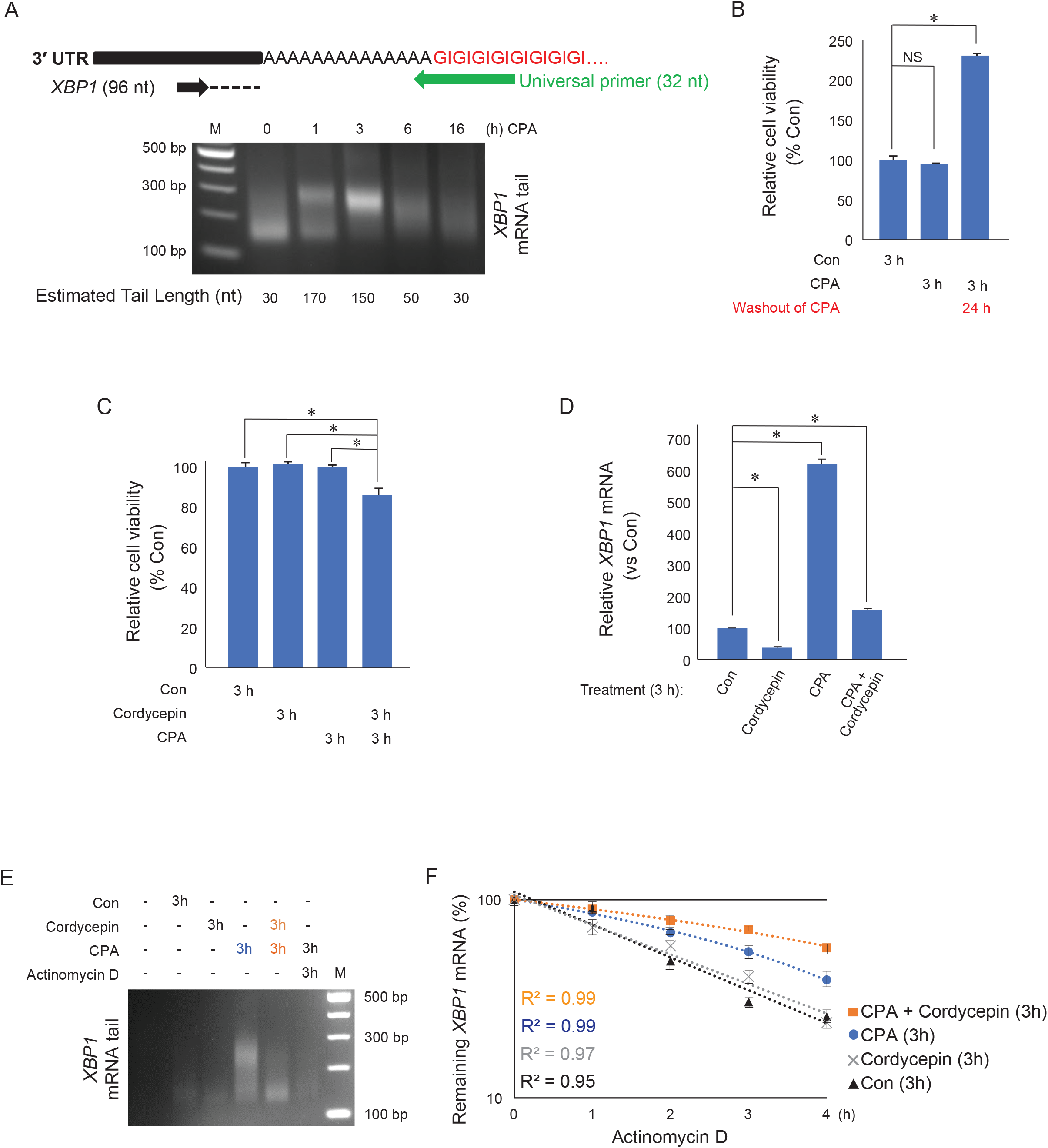
Nascent RNA polyadenylation during acute ER stress promotes cell survival. (A) MEFs were treated with CPA for the indicated durations. Following RNA isolation and the PCR-based poly(A) tailing assay, the poly(A) tail length of *XBP1* mRNA was measured in the indicated treatments. (B) MEFs were treated with DMSO (Con) or CPA (100 uM) for 3 h and cell viability was measured. Following CPA treatment, cells were allowed to recover in fresh media in the absence of CPA for 24 h past the 3 h treatment of CPA. (C) Cell survival was assayed in MEFs treated with DMSO (Con), CPA, 10 µg/ml Cordycepin, or combined CPA and Cordycepin for 3 h. (D) RT-qPCR analysis of *XBP1* mRNA levels in MEFs treated with DMSO (Con), CPA, Cordycepin, or combined treatment of CPA and Cordycepin for 3 h. (E) MEFs were treated with DMSO (Con), CPA, Cordycepin, or combined treatment of CPA with either Cordycepin or Actinomycin D for 3 h. *XBP1* mRNA poly(A) tail length was measured using the poly(A) tailing assay. (F) Cells were treated with DMSO (Con), CPA, Cordycepin, or combined treatment of CPA and Cordycepin for 3 h followed by Actinomycin D treatment for the indicated durations. Bars represent the mean of 3 measurements +/- SEM. The student’s t-test was used to evaluate the statistical significance; (*) P < 0.05. t1/2 = Con: 2 +/- 0.3 h, Cordycepin: 2.3 +/- 0.2 h, CPA: 3.5 +/- 0.3 h, and CPA+Cordycepin: 5.3 +/- 0.1 h

We next tested the effect of Cordycepin on *XBP1* mRNA stability during the acute UPR. Because Cordycepin inhibits accumulation of long poly(A) tail *XBP1* mRNA during the acute UPR (Fig. 4E), this experimental system allows us to determine if newly synthesized long poly(A) tail *XBP1* mRNA is critical for the observed longer half-life of the *XBP1* mRNA during the acute UPR. We found that the stability of the *XBP1* mRNA upon Cordycepin treatment together with CPA for 3 h was greater as compared to the stability of *XBP1* mRNA upon CPA-treatment alone (Fig. 4F). Furthermore, Cordycepin treatment in the absence of ER stress had no effect on the half-life of *XBP1* mRNA (Fig. 4F). These data suggest that the newly synthesized *XBP1* mRNA does not contribute to the observed increased *XBP1* mRNA stability during the acute UPR. Furthermore, these data support the notion that increased *XBP1* mRNA stability during the acute UPR reflects that of the short poly(A) tailed *XBP1* mRNA. Taken together, we propose that the acute UPR involves an unrecognized mechanism of selective mRNA translation via the synthesis of long poly(A) tail mRNAs that escape eIF2α-P-mediated translational repression and promote survival during the acute UPR and stabilization of pre-existing *XBP1* mRNA pool.

### ER stress-induced mRNAs have long poly(A) tails during the acute UPR

We next asked whether the temporal regulation of poly(A) tail length of *XBP1* mRNA was a feature of other UPR-regulated mRNAs. Based on published data, we selected the mRNAs for *ATF4* and *BiP* as UPR-induced genes and *SEC24D* and *ATP5B* as control genes not regulated by the UPR (Guan et al. 2017). We then evaluated the poly(A) tail length and abundance of the gene transcripts using the PCR-based tailing assay and RT-qPCR analysis, respectively, under UPR conditions. We found that the poly(A) tail length of *ATF4* and *BiP* mRNAs was long in response to the acute UPR, similar to the poly(A) tail length of the *XBP1* mRNA. In addition, the poly(A) tail length of these mRNAs had heterogeneous size in the chronic UPR (Fig. 5A). Furthermore, although the *ATF4* mRNA was induced in response to the UPR, its levels declined in the chronic UPR. On the other hand, *BiP* mRNA expression remained induced even following the transition to the chronic UPR (Fig. 5B). This latter can be explained by the long half-life of the *BiP* mRNA (Rutkowski et al. 2006). The non UPR-regulated *SEC24D* and *ATP5B* mRNAs did not show a significant change in their poly(A) tail length in response to the acute UPR; however, their poly(A) tail length was short during the chronic UPR, similar to the poly(A) tail of *GAPDH* mRNA (compare Figs. 2A and 5C). mRNA abundance for *SEC24D* and *ATP5B* showed negligible changes (Fig. 5D). We propose that in the acute UPR, newly synthesized mRNAs with long poly(A) tails are protected from translational repression, and the temporal regulation of poly(A) tail length is an important pro-survival mechanism of the UPR.

**Figure 5.**
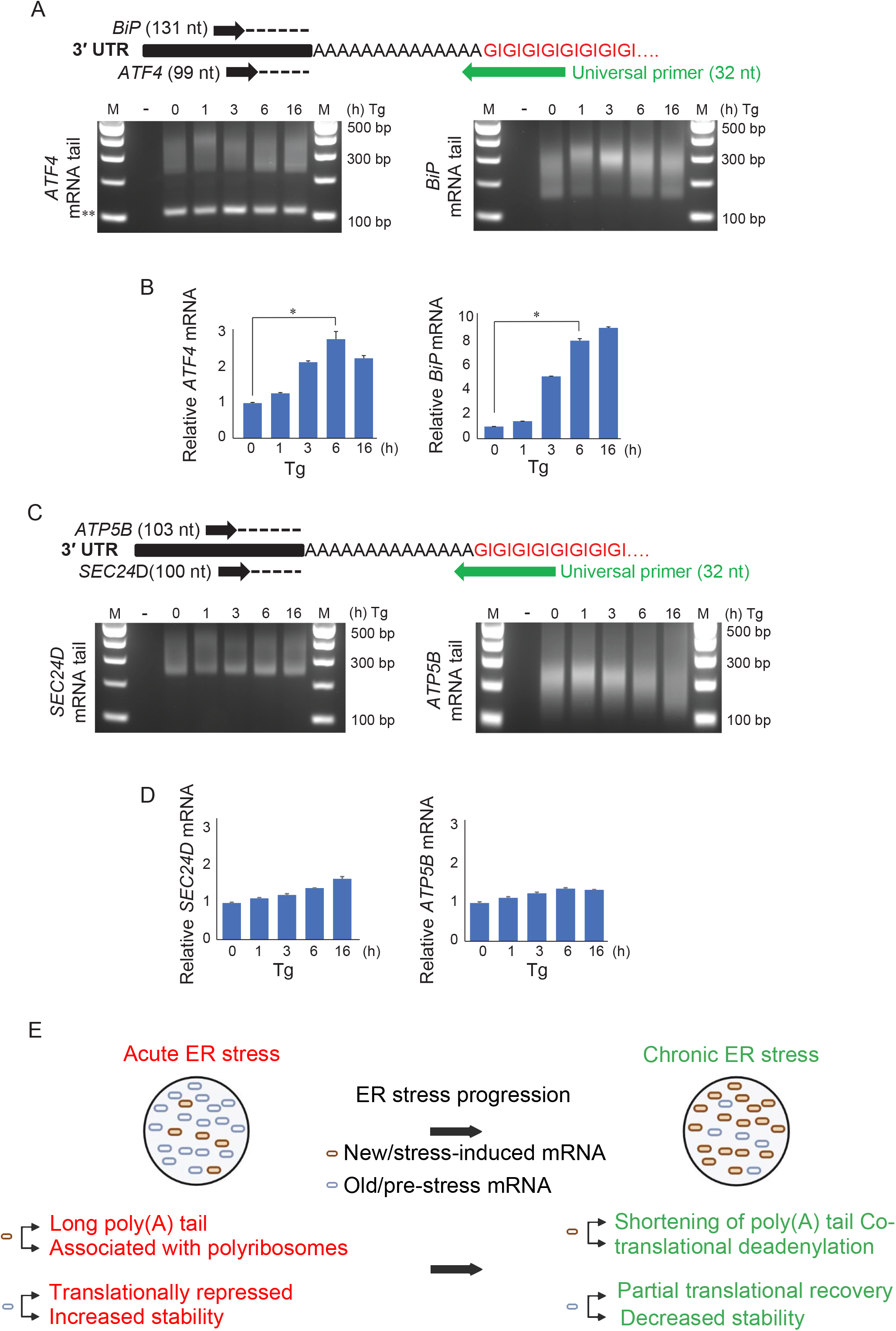
Temporal regulation of poly(A) tail length of UPR-regulated genes. (A) Experimental diagram of the PCR-based poly(A) tailing assay of *ATF4* and *BiP* mRNAs and their dynamic poly(A) tail length in MEFs treated with Tg for the indicated durations. The poly(A) tail length of *ATF4* and *BiP* mRNAs was estimated and a negative control (-) was used to monitor the tail reactions as previously explained. (**) indicates an artifact band. (B) The levels of *ATF4* and *BiP* mRNA were assessed using RT-qPCR in Tg-treated MEFs for the indicated durations. (C) Experimental diagram of poly(A) tailing assay of *SEC24D* and *ATP5B* mRNA and their dynamic poly(A) tail length in MEFs treated with Tg for the indicated times. (D) The level of *SEC24D* and *ATP5B* mRNA was assessed using RT-qPCR in Tg-treated MEFs for the indicated durations. (E) Model of *XBP1* mRNA regulation during acute and chronic ER stress. The steady state of the acute ER stress consists of different populations of *XBP1* mRNA. In the acute phase, the pre-ER stress (old) mRNA is translationally repressed and stabilized. However, the newly synthesized (new) mRNA has long poly(A) tails and escapes translational repression. In chronic ER stress, the pre-ER stress (old) mRNA is partially translationally derepressed and becomes unstable. Meanwhile, the newly synthesized (new) mRNA is subject to poly(A) tail shortening due to co-translational deadenylation. The figure was created by Biorender.com. Bars represent the mean of 3 measurements +/- SEM. The student’s t-test was used to evaluate the statistical significance; (*) P < 0.05.

## Discussion

We show here a new mechanism of translational control of *XBP1* gene expression during the UPR through the length of the poly(A) tail. During the acute UPR, we observed two discrete populations of *XBP1* mRNA, with either short or long poly(A) tails. The pool of long poly(A) tail-*XBP1* mRNA was derived from newly synthesized mRNA generated by stress-induced transcription of the *XBP1* gene and escaped translational inhibition. The pool of short poly(A) tail-*XBP1* mRNA, on the other hand, was translationally repressed and stabilized. Because the half-life of *XBP1* mRNA is short and its transcription is massively upregulated, progression from acute to chronic ER stress involves continuous enrichment with long poly(A) tail *XBP1* mRNA, while the mRNA with short poly(A) tails produced pre-stress is gradually depleted. Interestingly, during the chronic UPR, heterogeneous poly(A) tail length *XBP1* mRNA was translationally de-repressed and destabilized (Fig. 5E). Although the focus of this study was the regulation of the *XBP1* gene, we also showed similar biphasic regulation of poly(A) tail length of other stress-induced mRNAs, suggesting a more global mechanism of gene regulation during the UPR. We conclude that cells exhibit biphasic regulation of translation and stability during ER stress; in the acute phase, newly synthesized mRNA is protected from translational repression via the long poly(A) tails, thus initiating synthesis of pro-adaptive proteins. In the second phase of adaptation to chronic ER stress, expression of adaptive proteins is limited most likely via co-translational mRNA deadenylation and degradation. This biphasic response limits the UPR threshold and prolongs adaptation to chronic ER stress.

We observed here during the acute UPR the coupling of translation inhibition and increased stability of the *XBP1* mRNA with short poly(A) tails, while the newly synthesized *XBP1* mRNA with long poly(A) tails was insensitive to translational inhibition (Fig. 3). The phenomenon of coupling mRNA translation to stability was first described decades ago (Stimac et al. 1984; Peltz et al. 1992). Physiological processes such as mitosis or stress conditions that involve transient inhibition of protein synthesis are also characterized by transient mRNA stabilization (Ross 1997; Tanenbaum et al. 2015; Kawai et al. 2004; Majumder et al. 2012). The mechanisms for this regulation are still obscure; however, they are likely to involve the suggested loss of interaction between the stabilized mRNAs and factors of the degradation machinery when their translation is inhibited (Heck and Wilusz 2018; Morris et al. 2021). This would agree with more current reports of mRNA degradation occurring co-translationally (Pelechano et al. 2015). An alternative mechanism of regulation of mRNA stability via poly(A) tail length during the UPR may be via modification of the nucleotides of the poly(A) tail. Recent reports suggest that poly(A) tails may contain other nucleotides than adenosines, such as guanosines and uridines; guanylated poly(A) tails tend to be associated with long poly(A) tails and positively correlate with mRNA half-life, whereas uridylation is more common in short poly(A) tails and negatively correlates with mRNA half-life (Chang et al. 2014; Lim et al. 2018). It will be interesting to study if poly(A) tail modifications are involved in the biphasic mechanism of the regulation of poly(A) tail length during the UPR. For example, the assay shown in Fig. 2B could be used to identify if the different poly(A) tail lengths during the UPR consist of heterogeneous populations of modified poly(A) tails.

One striking observation in our study was the presence of two *XBP1* mRNA pools with different translational fates during the acute phase of the UPR; *XBP1* mRNA with short poly(A) tails was translationally attenuated as compared to newly synthesized pool harboring long poly(A) tails, which escaped translational inhibition. The mechanisms of generation of long poly(A) tail *XBP1* mRNA which escapes translational repression, was shown to involve transcriptional mechanisms during the UPR. However, it is unlikely to occur via regulation of polyadenylation because we have shown that the poly(A) tail length of newly synthesized *XBP1* mRNA is not significantly different between the unstressed, acute, and chronic UPR (Fig. 3D). The accumulation of long poly(A) tail *XBP1* mRNA parallels its increased synthesis during ER stress. Mechanisms linking transcription to mRNA translation, stability, and poly(A) tail length have recently been reported (Slobodin et al. 2020; Slobodin et al. 2017; Slobodin and Dikstein 2020). It was shown that reduced transcription dynamics correlated with enhanced m^6^A base modification, increased deadenylation activity, shorter poly(A) tails, and decreased mRNA stability of the target mRNAs (Slobodin et al. 2020). On the other hand, enhanced transcription correlated with less m^6^A deposition on mRNAs and positive regulation of translational efficiency (Slobodin et al. 2017). Such mechanisms can be investigated for the temporal regulation of *XBP1* mRNA poly(A) tail length during the UPR (Figs. 1C, D and 2A). Although most studies indicate weak correlations between translation efficiency and poly(A) tail length in somatic cells (Chang et al. 2014; Park et al. 2016; Subtelny et al. 2014), there is evidence that poly(A) tail length-mediated regulation of translation is likely dependent on the cellular context. For example, in zebrafish and frog embryos, poly(A) tail length is coupled to translation efficiency, but a developmental switch diminished this regulation during gastrulation (Subtelny et al. 2014). Here, we illustrated the preferential association of long poly(A) tail *XBP1* mRNA with heavy polyribosomes during the acute phase of the UPR (Fig. 3B). A possible explanation for this phenomenon might be that long poly(A) tail mRNAs are more competitive for the limited availability of poly(A) binding protein (PABP) (Xiang and Bartel 2021). Several studies have shown that PABP is a critical factor mediating translation initiation in mammalian cells (Martineau et al. 2008; Smith et al. 2017), and that during stress conditions that induce phosphorylation of the translation initiation factor eIF2α, PABP is sequestered in stress granules (Vanderweyde et al. 2013; Child et al. 2021). The latter can explain the limited PABP availability during the acute UPR, which can lead to competition among mRNAs and therefore selective mRNA translation. Systematic analysis of published data from genome-wide studies can determine poly(A) tail length requirements for the mRNAs under translational control during the acute UPR (Woo et al. 2018). We showed here that in the chronic UPR, most mRNAs tend to have short poly(A) tails (Fig. 2A, 5A, and 5C) at a time that their translation is de-repressed (Guan et al. 2017). This agrees with a previous report that short poly(A) tail mRNAs are characteristic of well-expressed genes (Park et al. 2016; Lima et al. 2017). However, our finding of decreased poly(A) tail length correlating with better translational efficiency during the chronic UPR seems contrary to the “dogma” that longer poly(A) tail mRNAs are better translated (Passmore and Tang 2021). If we consider our findings that shorter poly(A) tails of the *XBP1* mRNA were found on polyribosomes, the “dogma” may be correct, and the mechanism of generating shorter poly(A) tail mRNAs may involve co-translational deadenylation, as was shown previously (Webster et al. 2018; Vindry et al. 2012). Identifying the mechanism of co-translational deadenylation during the chronic UPR will suggest better experimental approaches to address the relationship between poly(A) tail length and efficiency of translation. Additional mechanisms for shortening of the poly(A) tails during the chronic phase could involve cis- and trans-regulatory elements such as miRNA and RNA-binding proteins recruited to mRNAs during ER stress, or release of PABP from stress granules (Backlund et al. 2020; Duan et al. 2020; Good and Stoffers 2020; Malhi 2014; Cairrao et al. 2021)

We described here the temporal regulation of poly(A) tail length of the newly synthesized *XBP1* mRNA during the UPR. A biphasic response to ER stress has previously been described by us and by others (Krokowski et al. 2013; Hetz and Papa 2018; Han et al. 2013). We show here that the *XBP1* gene is regulated in a temporal manner at the level of transcription, mRNA stability, and translation during acute and chronic ER stress (Majumder et al. 2012; Woo et al. 2018), further supporting the coordination of stress-induced gene regulation programs as an important mechanism of controlling the UPR threshold, which needs to be carefully controlled to maintain adaptation to chronic ER stress (Hetz and Papa 2018). Although partial recovery of the acute phase translational inhibition is necessary for adaptation to chronic ER stress (Guan et al. 2017), recovery of protein synthesis to normal pre-stress levels decreases adaptation to chronic ER stress via mechanisms involving the UPR-induced transcription program (Han et al. 2013; Krokowski et al. 2013). Furthermore, during chronic ER stress, selective translation initiation in the absence of active eIF4E involves the translation initiation factor eIF3D (Guan et al. 2017), which may be involved in the mechanism of ribosome-associated deadenylation of mRNAs during the chronic UPR. Here, we propose that the long poly(A) tails of newly synthesized mRNAs during the acute phase is a mechanism to escape translational repression, and the shortening of the poly(A) tails at the chronic phase may be a mechanism to limit the threshold of the UPR. The ribosome-associated shortening of poly(A) tails can contribute to both decreased translation efficiency and increased mRNA decay. Elegant studies by (Eisen et al. 2020) have shown that mRNA degradation occurs predominantly via deadenylation and identified that most mRNAs are committed to degradation once the poly(A) tail is 25 nucleotides or shorter. However, we cannot exclude the possibility that the gradient of poly(A) tail length during the chronic UPR also serves to decrease translational efficiency of mRNAs in subsequent rounds of translation initiation. Therefore, we propose that deadenylation during the chronic UPR can regulate either mRNA degradation or translation efficiency (Fig. 2B and 2C). This regulation of gene expression via poly(A) tail length is consistent with the adaptive response to chronic ER stress by keeping the UPR in homeostatic levels (Gomez and Rutkowski 2016; Guan et al. 2017).

The significance of the stabilization of the pre-existing *XBP1* mRNA in the acute phase of the UPR is not clear. Although it can be considered as a mechanism to amplify the chronic UPR response, an alternative function in the acute phase may be the anticipation of recovery from transient exposure to stress. mRNA stabilization during the acute phase may therefore preserve the capacity of the cells to synthesize proteins to facilitate faster recovery of ER function when exposure to stress is transient. This explanation is supported by other physiological responses such as cell cycle progression, where translation is inhibited transiently during mitosis and recovers as cells enter the G1 phase (Tanenbaum et al. 2015; Hume et al. 2020). Similar to the increased stability of translationally repressed mRNAs during the acute ER stress phase (Kawai et al. 2004; Majumder et al. 2012), translationally repressed mRNAs in mitosis are also stabilized (Ross 1997; Marzluff et al. 2008; Park et al. 2016).

In conclusion, our findings suggest an exciting hypothesis that establishing a steady state of adaptation to chronic ER stress involves a pre-steady state UPR phase of changes in transcription, translation, and mRNA stability. In the pre-steady state UPR, regulation of mRNA stability and translation promotes recovery from acute stress. In the steady-state adaption phase, the massive transcriptional induction and reprogramming of translation initiation protects cells during chronic ER stress. This biphasic response provides plasticity in the cellular response, allowing recovery from stress after acute or chronic episodes of ER stress with minimal commitment of resources and energy. Therefore, the coordinated transcriptional and post-transcriptional mechanisms of gene regulation in the pre-steady state and steady-state UPR is a critical cellular response to duration and intensity of environmental cues.

## Materials and Methods

### Cell Lines, Cell Culture Conditions, and Chemicals

Wild type (WT), IRE1α-deficient, and eIF2α-P-deficient mouse embryonic fibroblasts (MEFs) were grown in Dulbecco’s modified Eagle’s medium (DMEM) enriched with 10% fetal bovine serum (FBS) (Gibco), 100 units/mL penicillin, 100 µg/mL streptomycin, and 2 mM L-glutamine. Cells were grown at 37° C under 5% CO2. ER stress was induced by treating cells with either 400 nM Thapsigargin (Tg) (Sigma-Aldrich) or 100 µM Cyclopiazonic acid (CPA) (Tocris Bioscience) for the indicated durations. Other chemicals used in this study include 4µ8C (Torris Bioscience, 50 µM), Cycloheximide (CHX; 100 µg/mL) (Sigma-Aldrich), Puromycin (Puro; 5 µg/ml) (Thermo Fisher Scientific), Actinomycin D (ActD; 10 µg/ml), and Cordycepin (10 µg/ml) (Sigma-Aldrich).

### RNA Isolation, splicing reaction, and RT-qPCR

To measure mRNA levels and half-life, cells were seeded at 0.5 × 10^6^ cells in 60-mm culture dishes and grown to 80% confluency. Following the indicated treatments, total RNA was isolated using TRIzol (Invitrogen) according to the manufacturer’s instructions. The relative mRNA level was measured by the reverse transcription-quantitative polymerase chain reaction (RT-qPCR) normalized to levels of house-keeping genes such as *GAPDH*. Briefly, complementary DNA (cDNA) was synthesized using ProtoScript^®^ II Reverse Transcriptase with random primer mix (NEB). This cDNA was also used for a conventional PCR using Go-Taq^®^ Master Mix (Promega) with *XBP1* splicing primers. To illustrate whether *XBP1* mRNA was spliced, 2.5% ethidium bromide-stained agarose gels (Sigma) were prepared and run for 45 minutes at 150 volts. Also, mRNA abundance was quantitively determined using VeriQuest SYBR Green qPCR Master Mix (Affymetrix) with the StepOnePlus Real-Time PCR System (Applied Biosystems). Half-lives were calculated by fitting the data points to a nonlinear curve for the decay exponential of each target. Primers used in this study are listed in (Supplemental <u>Table S1</u>).

### RNA Labeling, Biotinylation, and Purification

WT MEF cells were seeded at 3 × 10^6^ cells in 150-mm culture dishes and were grown to 80% confluency. To metabolically label RNA, 400 µM 5-Ethynyl Uridine (5EU) (Click Chemistry Tools) was added for 1 h as indicated. Following total RNA isolation of labeled RNA with 5EU, we performed click chemistry using the CuAAC Biomolecule Reaction Buffer Kit-BTTAA based (Jena Biosciences). Following click chemistry, RNA was purified using spin columns (Zymo Research). RNA was resuspended in 50 µl nuclease-free water and mixed with equal volumes of 1X High Salt Wash Buffer (HSWB) (10 mM Tris pH 7.5, 1 mM EDTA pH 8, 0.1 M NaCl, 0.01% Tween-20). To capture the labeled RNA, 100 µl of Dynabeads(tm) MyOne(tm) Streptavidin T1 beads (Invitrogen) were used after being washed as described previously (Eisen et al. 2020). RNA from flow-through and elution was isolated using TRIzol LS (Invitrogen).

### PCR-based poly(A) tailing assay

Total RNA (10 µg) was incubated with recombinant RNAse-free DNase I (Sigma) in 50 µl-reaction tubes at 37° C for 20 minutes. The reaction was terminated by adding 200 µl of nuclease-free water and 750 µl of TRIzol LS (Invitrogen), and RNA was then isolated and requantified. To add an artificial tail to the mRNA poly(A) tail, 1.5 µg of the isolated RNA (up to 14.5 µl) was mixed with 2 µl of recombinant yeast Poly(A) Polymerase (PAP) together with 5 µl of 5X PAP buffer (Thermo Fisher Scientific), 2.5 µl of 10 µM GTP/ITP/MgCl2 (Jena Bioscience), and 1 µl of RNase inhibitor (New England Biolabs). The 25 µl-reaction was incubated in a thermocycler set at 37° C for 1 hour. G/I nucleotide-tailed RNA was then purified using spin columns (Zymo Research). The RNA, eluted in 9 µl, was used in a reverse transcription reaction as described previously (Patil et al. 2014) using the primer (CCCCCCCCCCTT) in the RT reaction. The resulting cDNA was used in a conventional PCR reaction to estimate the tail length. Primers used in this study are listed in (Supplemental <u>Table S2</u>). PCR products were run on 2.5 % ethidium bromide-stained agarose gels, and bands were visualized and captured under UV light (Syngene).

### RNA fractionation based on the length of poly(A) tails

The protocol used to visualize poly(A) tail lengths was based on the poly(A) tail fractionation performed by (Chorghade et al. 2017). Briefly, total RNA from untreated or Tg-treated cells for 3 or 16 h was isolated. 20 µl of total RNA (∼10-50 µg) was mixed with 200 µl of Guanidine Thiocyanate (GTC) buffer (4 M guanidine thiocyanate, 25 mM sodium citrate, pH 7.1), 4 µl of Beta-mercaptoethanol (β-ME), 5 µl of 50 pmol/µL biotinylated oligo-dT probes (Promega), and 408 µl of diffusion buffer (3X Saline-Sodium Citrate (SSC), 5 mM Tris, pH 7.5, 0.5 mM EDTA, 0.125% SDS, 5% ß-ME). The mixture was heated at 70° C for 5 minutes followed by a 10-minute centrifugation at 12,000 x g at room temperature. The supernatant was added to 150 µl of Dynabeads(tm) MyOne(tm) Streptavidin C1 beads (Invitrogen) that had been washed 3 times with 0.5X SSC buffer containing 0.02% Tween20. The beads were rotated slowly at RT for 15 minutes followed by 3 washes with 0.5X SSC buffer containing 0.02% Tween20. RNAs with the shortest poly(A) tails were eluted from the beads by adding 400 µl of 0.07X SSC followed by a 20-minute incubation at room temperature. The remaining fractions were eluted in the same way with the following SSC concentrations: 0.06X, 0.05X and 0.04X. The RNA from these fractions was purified by RNA-phenol/chloroform extraction, precipitated with sodium acetate and glycoblue (ThermoFisher) overnight, pelleted, and resuspended in 25 µL of water. 4 µL of RNA was 3’-end-labeled with ^32^pCp (prepared by incubating 16.5 µl of γ-^32^P-ATP (PerkinElmer), 1 µl of T4 polynucleotide Kinase, 2 µl of 10X buffer (NEB), and 833 µM of cytidine 3’-phosphate at 37° C for 1 hour, followed by 10 minutes at 65° C). The ^32^pCp was ligated using T4 RNA ligase 1 (NEB) and incubated overnight at 4° C. Following clean up with Micro Bio-Spin 6 columns (Bio-Rad), the 3’-end-labeled RNA was incubated with RNase A (Thermo Scientific(tm)) at 37° C for 30 minutes in a 100 µl-reaction containing 20 mM Tris pH 8, 1.25 mM MgCl2, 550 mM NaCl, 1.25 mM E. coli tRNA, and 0.125 mM RNase A. Radiolabeled poly(A) tails were purified by RNA-phenol/chloroform extraction and overnight precipitation with sodium acetate and glycoblue. The distribution of tail lengths was visualized by running the RNA on an 8.5% polyacrylamide gel with 7 M urea.

### Western Blot analysis

Cell extracts and western blot were performed as previously described (Guan et al. 2017). Briefly, Tg-treated MEFs were washed twice in cold PBS and lysed using a lysis buffer (50 mM Tris-HCl at pH 7.5, 150 mM NaCl, 2 mM EDTA, 1% NP-40, 0.1% SDS and 0.5% sodium deoxycholate) supplemented with EDTA-free protease inhibitor (Roche Applied Science) and PhosSTOP phosphatase inhibitor (Roche Applied Science). Cell lysates were placed on ice and sonicated 10 times. The supernatants of cold lysates were transferred to fresh tubes after centrifugation at 13,000 rpm for 10 minutes at 4 °C. The supernatants were used for protein quantification via the use of the DC(tm) Protein Assay Kit (Bio-Rad). Equal loading of proteins (10–20 µg) was analyzed in SDS-PAGE and primary antibodies were applied after a standard western blotting was performed. These antibodies were listed in (Supplemental <u>Table S3</u>).

### Metabolic Labeling of Cells with [^35^S] Methionine /Cysteine (Met/Cys)

eIF2α-P-deficient MEFs were seeded at 5 × 10^4^ cells/well in 24-well plates. Cells were grown overnight and treated as described above. Prior to the end of each treatment, [^35^S]-Met/Cys was added (30 μCi/mL EXPRE^35^S Protein Labeling Mix, PerkinElmer) for 30 minutes. Next, cells were washed twice with cold phosphate-buffer saline (PBS) and total proteins were precipitated in 5% trichloroacetic acid (TCA) with 1 mM Methionine (Met) (Sigma-Aldrich) for 15 minutes on ice. The precipitation step was repeated overnight at 4°C. After careful removal of TCA-Met, 200 μL of 1 N NaOH and 0.5% sodium deoxycholate were added for 30 minutes. To determine the incorporation of [^35^S]-Met/Cys into total cellular proteins, liquid scintillation counting, and DC Protein Assay (Bio-Rad) were used to quantify radioactivity and protein concentration, respectively.

### Polysome Profile Analysis and PCR-based tailing assay

Wild-type MEF cells were seeded at 3 × 10^6^ cells/150-mm culture dishes (2 dishes per treatment) and grown to reach 80% confluency. Cells were washed twice with cold PBS containing 100 μg/ml cycloheximide and placed on ice. 1 ml of lysis buffer (10 mM HEPES-KOH (pH 7.4), 5 mM MgCl2, 100 mM KCl, 1% Triton, 100 μg/ml cycloheximide, 2 mM DTT, 200 units/ml RNase inhibitor (New England Biolabs), EDTA-free protease inhibitor (Roche Applied Science)) was added to each plate after removing the remaining PBS carefully. Next, cells were scraped, and then passed 5 times through a 26-gauge needle. Lysates were spun at 13,000 x g for 15 minutes, and supernatants containing cytosolic cell extracts were collected. Approximately 10 A units (260 nm) of lysates were layered over 10%–50% cold sucrose gradients in buffer (10 mM HEPES-KOH (pH 7.4), 2.5 mM MgCl2, 100 mM KCl). Gradients were centrifuged at 31,000 rpm in a Beckman SW32 rotor for 2.5 h at 4° C. After centrifugation, gradients were fractionated and collected into 12 tubes (∼1 ml/fraction). RNA from each fraction was isolated using TRIzol LS (Invitrogen), and an equal volume of RNA from each fraction was used for cDNA synthesis. The relative quantities of specific mRNAs were measured by quantitative RT-PCR (RT-qPCR) as described above. To measure the poly(A) tail length of mRNAs, fractions were equally combined into 3 pools of free, light, and heavy polyribosomes. After RNA isolation, tail length was measured using equal volumes from each pool of mRNAs as described above.

### Cell Viability

WT MEF cells were seeded at 1 × 10^4^ cells/well in 96-well plates. Cells were grown overnight and treated as indicated. Prior to the end of treatment, an equal volume of CellTiter-Glo® reagent (Promega) was added to the existing media volume. After mixing the reagent well, the plate was incubated at 37° C for 10 minutes, followed by incubation at room temperature on a shaker for another 10 minutes. Luminescence was recorded using a SpectraMAx M3 instrument.

### Statistical Analysis

The mean of triplicate measurements was statistically evaluated using the student’s t-test, where (*) indicates significance with P < 0.05. Otherwise, results were considered non-significant (NS). Error bars indicate standard error of the mean.

## Supporting information

all supplemental data for Alzahrani (2022)

## Acknowledgments

This study was conducted with the support of National Institutes of Health grants (R01DK53307, R01DK060596, and R01DK113196) (MH) and CDDRCC pilot grant DK097948 (MH), GM143364 (KEB) and GM107331 (DDL). We thank Dr. Randal Kaufman for providing cell lines of WT-MEFs and eIF2α-P-deficient MEFs and IRE1α-deficient MEFs. We, also, thank Dr. Raul Jobava for his assistance with the polysome profile analysis in this manuscript.

