## Supplementary material for "Newly synthesized mRNA escapes translational repression during the acute phase of the mammalian unfolded protein response": all supplemental data for Alzahrani (2022): Supplemental_Figure legends.pdf.pdf

**Supplemental Figure S1**

**Evaluation of *XBP1* mRNA splicing and turnover in different cell lines using RT-qPCR**. (A) (top) RT-PCR analysis of *XBP1u* mRNA splicing in IRE1α-deficient and WT MEFs, treated with Tg for 0, 1, and 16 h, or eIF2α-P-deficient MEFs treated with Tg and 4μ8C together for the indicated durations. (bottom) Splicing efficiency of *XBP1u* mRNA in the indicated cell treatments evaluated by RT-qPCR analysis. (B) The half-life of the *XBP1u* mRNA was measured in the indicated cell line and treatments. Harringtonine, a translational initiation inhibitor was used at 2 μg/ml for 1 h. Bars represent the mean of 3 independent determinations +/- SEM.

**Supplemental Figure S2**

**Distribution of mRNAs on polysome profiles as a measurement of their translation efficiency**. (A-B) Polysome profile distribution of *XBP1u* and *ATF4* mRNAs in IRE1α-deficient MEFs treated with Tg for 0, 1, and 16 h in cell extracts analyzed on sucrose gradients (10% to 50%). The enrichment of these mRNAs in the last 3 fractions of each condition was evaluated. (C) Distribution of *XBP1u* mRNA in polysome profiles (as in A) of eIF2α-P-deficient MEFs treated with Tg for 0, 1, and 16 h in the presence of 4μ8C. The enrichment of *XBP1u* mRNA in the last 3 fractions of each condition was evaluated.
