## Supplementary material for "Newly synthesized mRNA escapes translational repression during the acute phase of the mammalian unfolded protein response": all supplemental data for Alzahrani (2022): Supplemental_Table_S1.pdf.pdf

**Supplemental Table S1: List of primers used in RT-qPCR**

| Target/ Primers | Forward primer | Reverse Primer |
|---|---|---|
| *GAPDH* | CGCCTGGAGAAACCTGCCAAGTATG | GGTGGAAGAATGGGAGTTGCTGTTG |
| *XBP1s* | GAGTCCGCAGCAGGTG | CTGGGAGTTCCTCCAGACTA |
| *XBP1u* | GACTATGTGCACCTCTGCAG | CTGGGAGTTCCTCCAGACTA |
| *XBP1* total | GGCTGTCTGGCCTTAGAAGA | CTGTCAAATGACCCTCCCTG |
| *XBP1* Splicing* | ACACGCTTGGGAATGGACAC | CCATGGGAAGATGTTCTGGG |
| *Sec24D* | AGCCTGAAATCTGTCTGGTAGA | CCGTTTTATAGACAAAACAACTGG |
| *ATP5B* | GATGTGATGTTCTCTCTGAAGAG | CCACCACTGTGAGCTCAA |
| *HSPA5/BiP* | AGGGTGTGTGTTCACCTTGG | AACATTTATTGGTGTCACTTATGGT |
| *ATF4* | GAGGCTCTGAAAGAGAAGGCAG | CAAGCACAAAGCACCTGACTAC |

* Primers used for RT-PCR.
