## Supplementary material for "Newly synthesized mRNA escapes translational repression during the acute phase of the mammalian unfolded protein response": all supplemental data for Alzahrani (2022): Supplemental_Table_S2.pdf.pdf

**Supplemental Table S2: List of primers used in the poly(A) tailing assay**

| Target/ Primers | Forward primer | Reverse Primer |
|---|---|---|
| GI-Tail | | GAGTAGCGTTGAATAAGTTGCCCCCC CCCCTT |
| *GAPDH* | ACTGAGCAAGAGAGGCCCTATC | GTTATTATGGGGGTCTGGGATGG |
| *XBP1* | GTAAATGCTTGATGGATCTTCTTGC | GCTGTGTTGCTTTTTTTTTAATTGC |
| *Sec24D* | AGCCTGAAATCTGTCTGGTAGA | CCGTTTTATAGACAAAACAACTGG |
| *HSPA5/BiP* | AGGGTGTGTGTTCACCTTGG | AACATTTATTGGTGTCACTTATGGT |
| *ATF4* | GAGGCTCTGAAAGAGAAGGCAG | CAAGCACAAAGCACCTGACTAC |
| *ATP5B* | GATGTGATGTTCTCTCTGAAGAG | CCACCACTGTGAGCTCAA |
