## Supplementary material for "Newly synthesized mRNA escapes translational repression during the acute phase of the mammalian unfolded protein response": all supplemental data for Alzahrani (2022): Supplemental_Table_S3.pdf.pdf

**Supplemental Table S3: List of primary antibodies used in western blotting analysis**

| Antibodies | Company | Catalog number |
|---|---|---|
| Rabbit monoclonal anti-ATF4 | Cell Signaling Technology | #11815 |
| Rabbit monoclonal anti-BiP | Cell Signaling Technology | #3177 |
| Rabbit polyclonal anti-eIF2α | Cell Signaling Technology | #9722 |
| Rabbit monoclonal anti-eIF2α-P (Phosphorylated at Ser 51) | abcam | #ab32157 |
| Mouse monoclonal anti-α-tubulin | Sigma-Aldrich | #T9026 |
| Rabbit polyclonal anti-XBP1s | Cell Signaling Technology | #83418 |
