## Supplementary figures and images for "Newly synthesized mRNA escapes translational repression during the acute phase of the mammalian unfolded protein response"

### Supplemental_Fig_S1.pdf.pdf

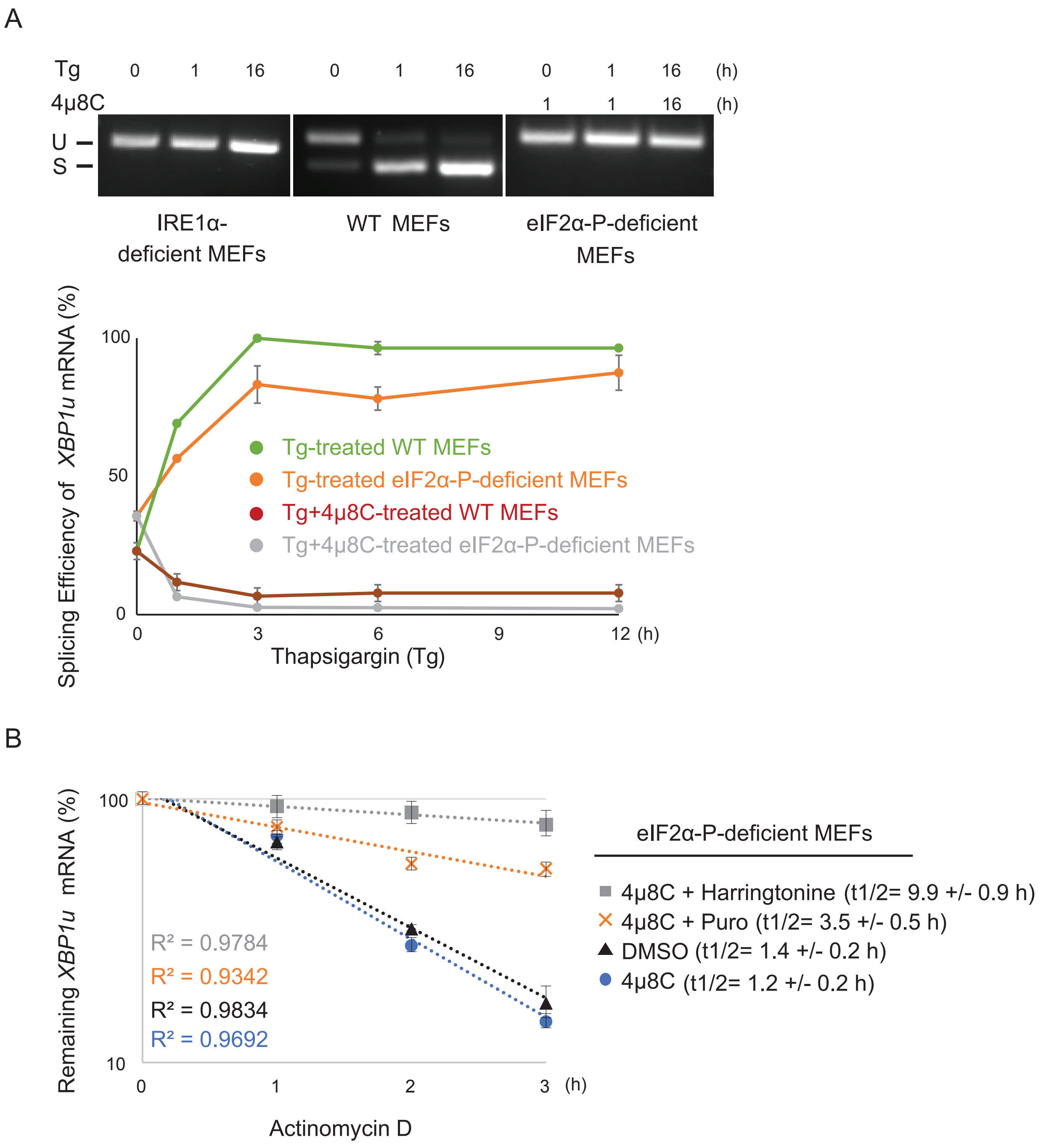

### Supplemental_Fig_S2.pdf.pdf

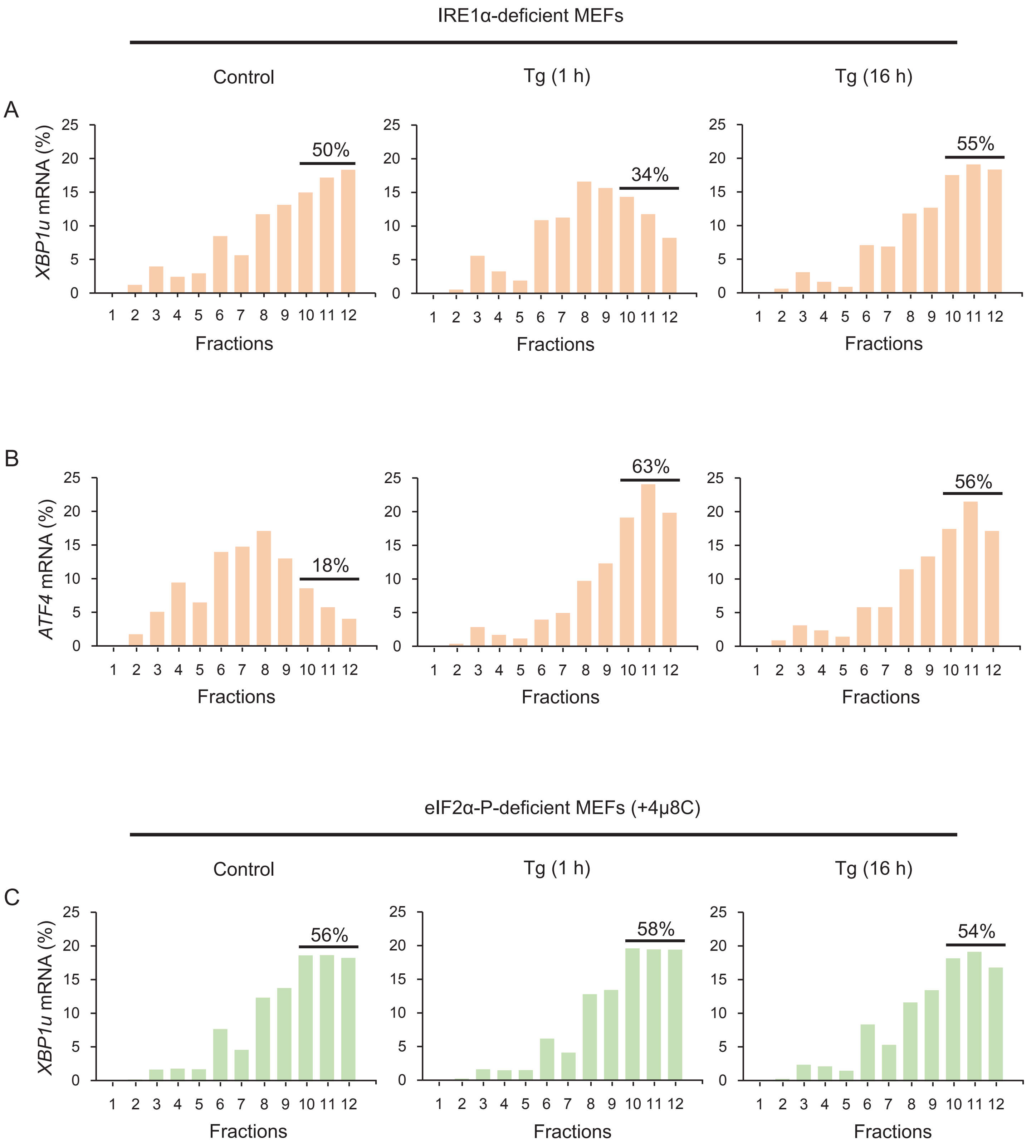


Supplemental_Fig_S2 Alzahrani et al. (2022)
